# Intrinsic control of muscle attachment sites matching

**DOI:** 10.1101/544569

**Authors:** Alexandre Carayon, Laetitia Bataillé, Gaëlle Lebreton, Laurence Dubois, Aurore Pelletier, Yannick Carrier, Antoine Wystrach, Alain Vincent, Jean-Louis Frendo

## Abstract

How a stereotypic muscle pattern is established, and adapted to fit locomotion behaviour is a fascinating question. Here we set up the targeted deletion of cis-regulatory modules (CRMs) controlling the transcription of *Drosophila* muscle identity transcription factors (iTF) to generate larval muscle identity mutants. By focusing on one muscle transcription and morphology, we show that selection of muscle attachment sites and the precision of muscle/muscle matching is intrinsic to muscle identity. It involves propagation of the iTF expression code from a founder myoblast to the other syncytial nuclei after fusion. Live imaging indicates that the precise staggered muscle attachment pattern involves attraction to tendon cells and homotypic repulsion. Unbalance leads to the formation of abnormal, branched muscles. Single muscle morphology shifts induce subtle locomotor behaviour. Together this work highlights that CRM deletion is an effective setting for creating muscle-specific defects and branched muscles, as new paradigms to study the development of human myopathies affecting subsets of muscles.

**Highlights:**

– First muscle-identity mutants, via deletion of specific cis-regulatory modules
– Reprogramming of syncytial nuclei is key to muscle morphological identity
– Selection of muscle attachment sites; attraction and retraction intrinsic to muscle identity
– Genetically controlled formation of branched muscles, a new paradigm for functional studies
– Single muscle morphology shift induces subtle locomotor behaviour modification

## Background

The musculature of each animal is composed of a complex array of body wall muscles allowing precision and stereotypy of movements. The somatic musculature of the *Drosophila* larva - about 30 distinct body wall muscles per each hemi-segment, organized into three layers, internal, median and external - is a suitable model to study how a muscle pattern is specified and linked to locomotion behaviour. Each muscle is a single multinucleated fibre with a specific morphological identity - size, shape, position along the dorso-ventral (DV) and antero-posterior (AP) axes. The muscles are attached to the exoskeleton via tendon cells at precise positions along segmental borders and within a segment [1–3]. Internal muscles such as the dorsal acute (DA) muscles, adhere both to segmental tendon cells and a muscle(s) in the next segment, forming “indirect” Muscle Attachment sites ([4], for review), adding complexity to the 3-D architecture of the musculature.

*Drosophila* muscle development proceeds through fusion of a founder myoblast (founder cell, FC) with fusion-competent myoblasts (FCMs) ([5], for review). FCs originate from asymmetric division of progenitor cells (PCs) which are themselves selected from equivalence groups of myoblasts, called promuscular clusters (PMCs). Muscle identity ensues from the expression by each PC and FC of a specific combination of identity transcription factors (iTFs) [6], established in three steps: First, different iTFs are activated in different PMCs, in response to positional information, and this expression is only maintained in PCs. Second, refinement of the FC iTF code occurs via cross-regulations between iTFs in PCs and/or FCs and Notch signalling [7–9]. Several PCs can be serially selected from a PMC and give rise to different muscle identities according to birth order, adding a temporal dimension to the positional regulation ([10]; Figure 1 Sup1). A well-documented example is the distinction between two DA identities, DA2 and DA3 [11]. A third step is the transcriptional activation of iTFs in other syncytial nuclei after fusion [12, 13] which correlates with the progressive activation of identity “realisation” genes acting downstream of iTFs [14, 15]. Some iTFs are expressed transiently, however, other at every step during muscle development and the relative contributions of PC/FC *versus* syncytial expression to conferring a muscle its final morphology remain unclear. The consequence of specific muscle identity defects on locomotion, a question of prime importance for progress on studying human myopathies which affect subsets of muscles, remains largely to be assessed.

Genetically controlled muscle identity changes should provide suitable models for studying locomotion deficits linked to muscle morphology imbalance. However, mutations for known iTFs are embryonic lethal and/or show pleiotropic phenotypes reflecting their multiple expression sites. Here, we took advantage of our previous characterization of Col expression in a single larval muscle, the dorso-lateral DA3 muscle. This expression is under control of two sequentially acting *col* cis-regulatory modules (CRMs), which we thought could be used to generate muscle-specific mutants and circumvent lethality/pleiotropy problems.

CRM deletion results show the early (*col*-ECRM), and late (*col*-LCRM) CRMs are redundant at the PC stage, emphasizing that it is a key step in specification of muscle identity. Full *col*-LCRM deletion results into loss of *col* transcription in DA3 FCs and DA3 into DA2 morphological transformation. Removal of the auto-regulatory module alone prevents col activation in syncytial nuclei and leads to incomplete transformations, *i.e.*, branched DA2-DA3 muscles. Together, it shows that i) FC identity must be propagated to syncytial nuclei for a muscle to adopt a specific morphology; ii) specific staggered muscle/muscle attachments involve homotypic repulsion; iii) branched DA2-DA3 muscles affect larval locomotion performance. Branched muscles are typical of late, severe phases of human Duchenne Muscular Dystrophy. Drosophila CRM deletion is an effective setting for creating muscle-specific defects and branched muscles, as many paradigms to study myopathies and pathological muscle regeneration in humans.

## Results

### Redundant CRMs ensure robust PC-specific iTF transcription

With the prospect of using CRM deletions to generate muscle-specific mutants, we re-assessed *col* regulation in the DA3 muscle lineage. At embryonic stage 10, *col* transcription is expressed in a dorso-lateral PMC which gives rise to 3 PCs, DA2/AMP, DA3/DO5 and LL1/DO4, born in this temporal order, in thoracic and abdominal segments, and the DT1/DO3 PC in abdominal segments [10] (Figure 1 Sup1). It is maintained in the DA3/DO5 and DT1/DO3 PCs at stage 11 (Figure 1 Sup1), before being restricted to the DA3 FC at stage 12 and activated in FCM nuclei recruited into the DA3 growing fiber between stages 13 to 16 [12]. Two CRMs together reproduce this dynamics of expression in reporter assays: *col*-ECRM (previously CRM276; [9]) is active in the PMC and transiently the DA3/DO5 and DT1/DO3 PCs, *col*-LCRM in the same PCs and the DA3 FC and syncytium [16] (Figure 1) *col*-ECRM was identified by a computational search for cluster of transcription factor binding sites, and *col*-LCRM by functional dissection of the *col* upstream region in reporter assays [9, 16]; (Figure 1) and both overlap in *vivo* binding sites for the master myogenic TFs, Mef-2 and Twist (Twi) [17, 18] (Figure 1A-B). This did not exclude the possible existence of redundant enhancers [19]. We therefore conducted a systematic analysis of Gal4 reporter lines [20, 21] covering 36kb of the *col* genomic region (Figure 1A). Only one line, GMR13B08, reproduced PMC Col expression and it overlaps *col*-ECRM. Likewise, only the GMR12G07 and 12H01lines which partly overlap *col*-LCRM displayed DA3 expression (data not shown, Figure 1A). We could thus conclude to the absence of *col* mesodermal CRM other than *col*-ECRM and *col*-LCRM.

**Figure 1.**
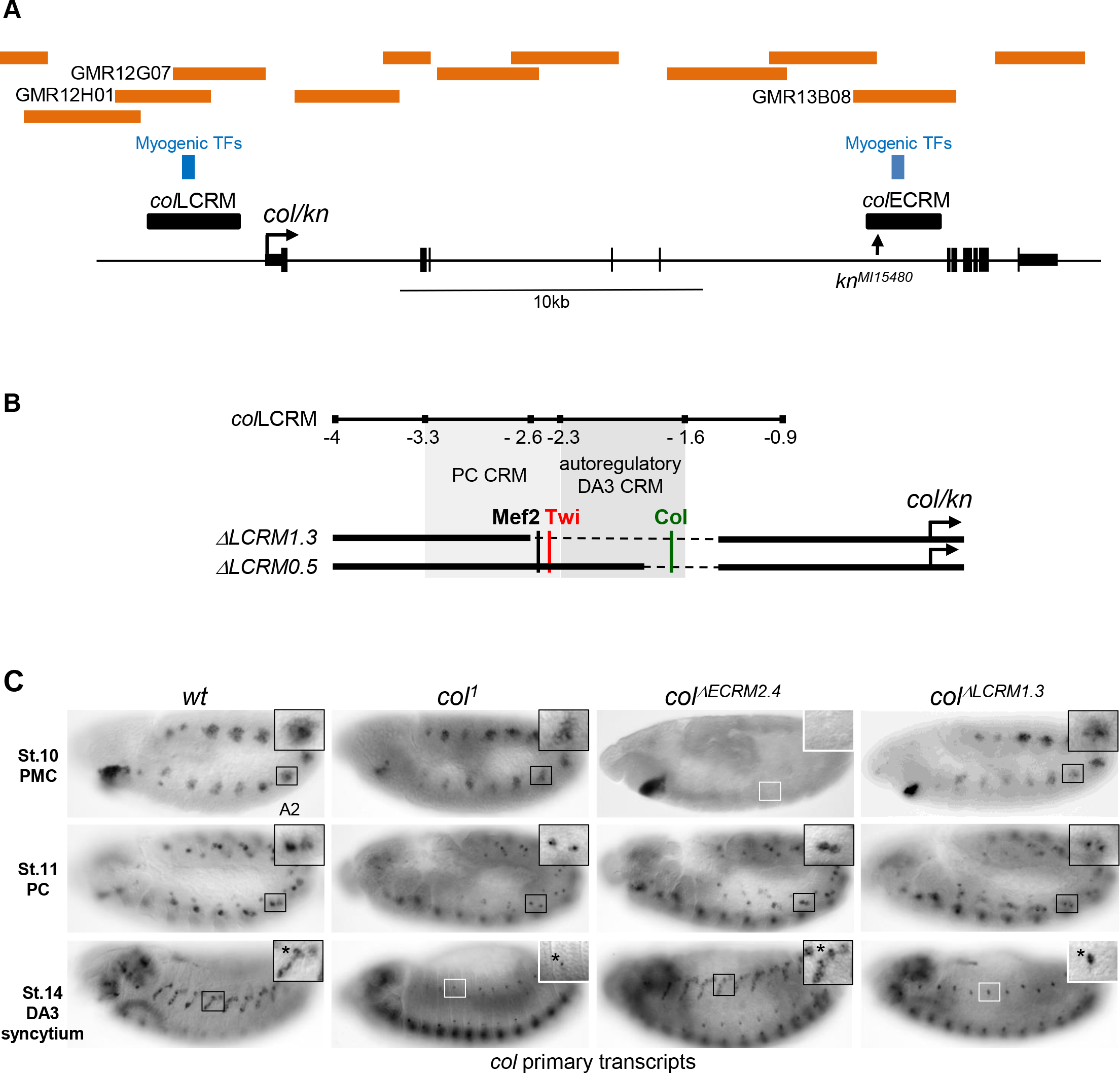
*col* CRMs, CRM deletions and *col* transcription. **(A)** Schematic representation of the *col* transcribed region, transcript A (http://flybase.org/cgi-bin/gbrowse2/dmel/?Search=1;name=FB_gn0001319). GMR and VT fragments tested in reporter assays are indicated by brown horizontal bars; only those numbered were found active in muscles. They overlap the previously mapped early *col*-ECRM and late *col*-LCRM [9] (black bars). The positions of clusters of binding sites for the myogenic TFs Mef-2, Twi and Tin [17, 60] are shown by vertical blue bars, and the *kn*^*Mi15480*^ transposon insertion by a vertical arrow. **(B)** Enlarged view of *col*-LCRM indicating the positions of juxtaposed PC-specific and autoregulatory DA3-specific CRMs, Col, Mef2 and Twi binding sites, and the *col*^*ΔLCRM1.3*^ and *col*^*ΔLCRM0.5*^ deletions generated by CRISPR/cas9 genome editing. **(C)***col* transcription in wt and mutant embryos, genotypes indicated on top, visualised by *in situ* hybridisation to primary transcripts. A detail of the abdominal A2 segment (squared area) is shown in the top right corner in each panel. Stage 10 embryos, PMC *col* transcription is lost in *col* ^*ΔECRM2.4*^ embryos (white square); stage 11, *col* transcription in the DA3/DO5 and DT1/DO4 PCs is detected in all strains; stage 14, DA3 syncytium transcription is lost in *col*^*1*^ and *col* ^*ΔLCRM1.3*^ embryos (white squares). *indicates col transcription in a multidendritic (md) neuron.

Based on this conclusion, we separately deleted from the genome a 2,4kb fragment removing *col*-ECRM (Figure 2 Sup1), a 1.3kb fragment removing the core region of *col*-*LCRM* [16] and a 0.5kb *col*-*LCRM* fragment removing the Col autoregulation site [22] (Figure 1B; Figure 2 Sup2). The resulting *Drosophila* strains are referred to as *col* ^*ΔECRM2.4*^, *col* ^*ΔLCRM1.3*^ and *col* ^*ΔLCRM0.5*^, respectively (Figure 1B). Both *col* ^*ΔLCRM*^ strains are homozygous viable and fertile. *col* ^*ΔECRM2.4*^ strains are homozygous female sterile, a sterility unrelated to *col* activity, since fertility is restored by placing *col*^*ΔECRM*^ over a deficiency (*Df(2L)BSC429*) removing the entire *col* locus (data not shown). As a first step to determine the myogenic functions of *col*-ECRM and *col*-LCRM in muscles, we compared *col* transcription between wt (+/+), *col*^1^ (a protein null mutant), *col*^*ΔECRM2.4*^ and *col* ^*ΔLCRM1.3*^ strains, using an intronic probe to detect nascent transcripts (Figure 1C). In *col^1^* homozygous mutant embryos, *col* transcription is detected in PMC cells as in wt, weaker in PCs, and undetectable at the FC stage, indicating the key role of autoregulation from that stage [12, 22] (Figure 1C). In *col* ^*ΔECRM2.4*^ embryos, *col* is not activated in PMC cells, as could be expected. Yet, low *col* transcript level is detected in the DA3/DO5 and DT1/DO3 PCs. This shows that inheritance of Col protein synthesized under *col*-ECRM control is not required for activation of *col* transcription in these PCs. Accordingly, a normal pattern is then observed in the developing DA3 muscle, stage 14. Conversely, *col* is transcribed in PMC cells and the DA3/DO5 and DT1/DO3 PCs in *col* ^*ΔLCRM1.3*^embryos, reflecting *col*-ECRM activity (see above), but not in the developing DA3 muscle, showing that *col*-LCRM is strictly required for this expression.

*col*-LCRM activity in PCs in absence of Col inheritance is different from *col* autoregulation previously mapped to the −2.3 to −1.6 *col* upstream region [16, 22] (Figure 1B; Figure 1 Sup2). Analysis of additional *lacZ*^*yi*^ reporter genes designed to detect primary transcription [9, 23] showed that the −3.3 to −2.3 fragment is only active in PCs, thus defining a PC-specific CRM (Figure 1 Sup2C; Figure 1B). This CRM contains conserved *in vivo* binding sites for Mef2 and Twist, diagnostic of mesoderm/somatic muscle CRMs [18] (Figure 1A; Figure 1 Sup2B). Deletion of a 97bp fragment removing both sites indeed abolishes reporter transcription (Figure 1 Sup2). We conclude that *col*-LCRM is in fact the juxtaposition of a PC-specific CRM containing Mef2 and Twi binding sites and a DA3-specific maintenance CRM mediating *col* autoregulation (Figure 1B). One unexpected outcome from the preliminary characterization of *col*-ECRM and *col*-LCRM deletions was their functional redundancy in two specific PCs. This redundancy emphasizes that the PC stage is when robust iTF expression is critical to muscle identity specification.

### *col* CRM deletions result in muscle transformation and branched muscles

The redundant activity of *col*-ECRM and *col*-LCRM in PCs allowed us to assess the relative contribution of PC *versus* FC-muscle expression in specifying muscle identity. Because of additional unknown mutation in the *col* ^*ΔECRM2.4*^ line, responsible on female sterility, and to avoid any ectopic effect, we therefore placed each wt and CRM deletion chromosome over the deficiency *Df(2L)*^*BSC429*^ chromosome (abbreviated *Df* in some Figure ures) which lacks the entire *col* locus. The *col*-LCRM-moeGFP reporter is used to visualize DA3 morphology [24]. Control +/*Df(2L)*^*BSC429*^ hemizygous embryos show a wt pattern of *col*-LCRM-moeGFP expression, *i.e*, expression in the DA3/DO5 and DT1/DO3 PCs at stage 11 and the DA3 muscle at stage 15 (Figure 2A). *col*-LCRM-moeGFP expression in *col ^*ΔECRM2.4*^/Df(2L)^BSC429^* is similar to wt and the DA3 morphology is affected in only 20% of segments (Figure 2A-B). On the contrary, it is barely detected in *col*^*ΔLCRM1.3*^embryos at stage 15, consistent with ISH data (Figure 1C). Residual GFP expression showed that the DA3 muscle is most often (85.2% of segments) transformed into a more longitudinal, DA2-like muscle (designated below as DA3^>DA2^; Figure 2A-B), similar to observed in *col* null mutant embryos [24]. The DA3^>DA2^ morphology reflects its anterior attachment to dorsal, rather than lateral tendon cells. Of interest, a number (6.3%) of branched muscles corresponding to three attachment sites, two anterior and one posterior, are also observed (Figure 2A). Branched (DA3^>DA3/DA2^) muscles are observed in higher proportions (29%) than DA3^>DA2^ transformations in *col ^*ΔLCRM0.5*^/Df(2L)^BSC429^*, *i.e*, when only the autoregulation module has been deleted (Figure 2A-B). In view of the low numbers of DA3 transformations in *col* ^*ΔECRM*^ embryos and high ratios of complete or incomplete DA3 transformations in *col* ^*ΔLCRM*^ deletion mutants, we pursued the analysis of *col*^*ΔLCRM1.3*^ and *col*^*ΔLCRM0.5*^ mutants. Both phenotypes show that iTF CRM deletion strategy is effective for creating viable muscle-specific identity mutants.

**Figure 2.**
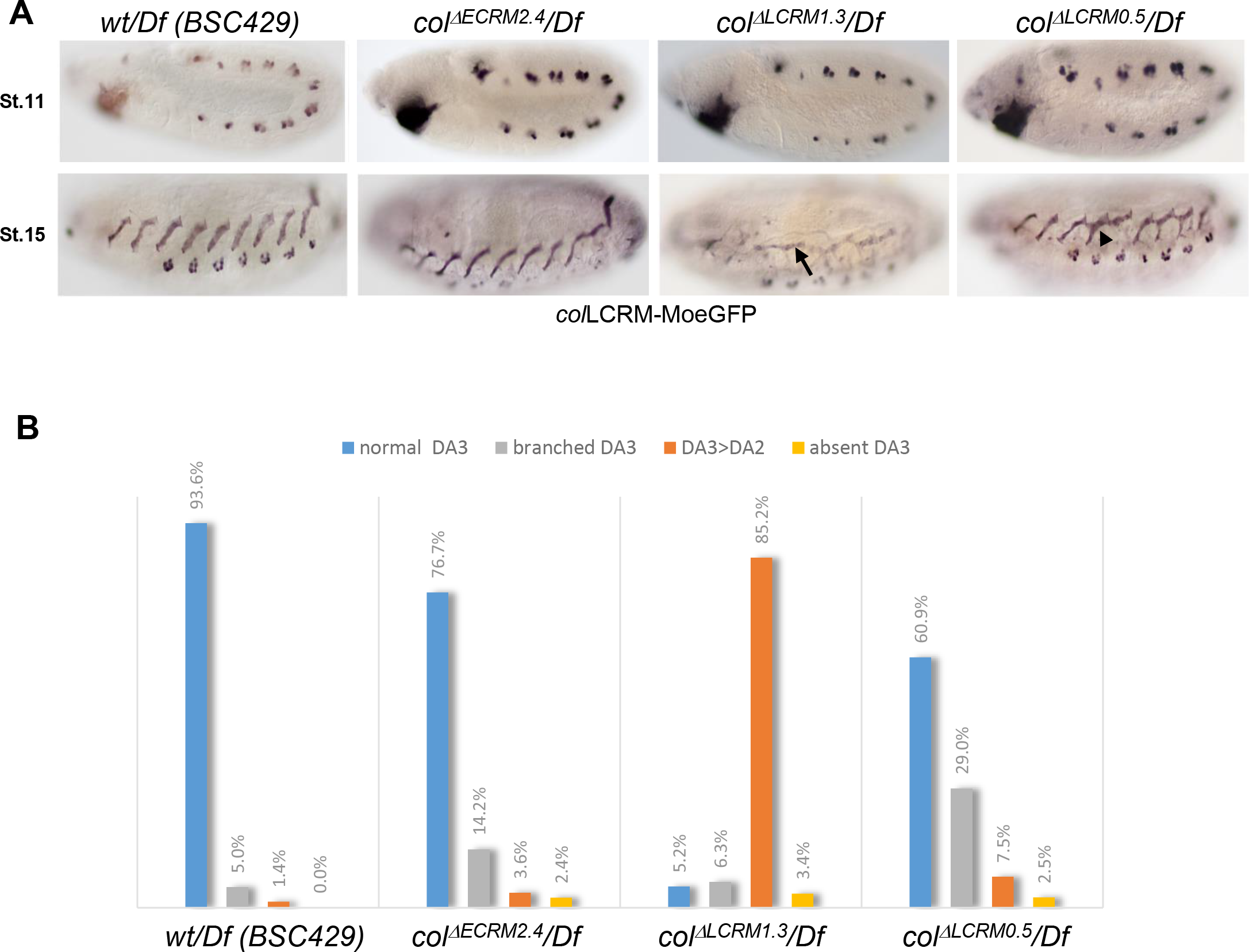
Muscle transformations upon *col*-*CRM* deletions. **(A)** *col*-LCRM-moeGFP expression in stage 12 and 15 embryos in hemizygous *Df(2L)^BSC429^*; *col*-LCRM-moeGFP (abbreviated *Df*)/*wt*, *col* ^*ΔECRM2.4*^, *col* ^*ΔLCRM1.3*^ and *col* ^*ΔECRM0.5*^ embryos, as indicated on top. GFP expression in PCs at stage 11 is similar in all strains. DA3^>DA2^ transformations (arrow) and branched DA3-DA2 (DA3^>DA3/DA2^) muscles (arrowhead) are observed in *col* ^*ΔLCRM*^ embryos. **(C)** Quantification of the relative proportions of DA3, DA3^>DA2^, branched DA3-DA2 and absence of DA3 muscles in *Df*/*wt*, *col* ^*ΔECRM2.4*^, *col* ^*ΔLCRM1.3*^and *col* ^*ΔECRM0.5*^ embryos.

### Reprogramming of syncytial nuclei is required for specific muscle morphology

The complete DA3^>DA2^ and incomplete DA3^>DA3/DA2^ identity shifts observed in *col* ^*ΔLCRM1.3*^ and *col* ^*ΔLCRM0.5*^, respectively (Figure 2), raised the possibility that reprogramming of fused FCM nuclei differed between these two strains (Figure 3). Here, (re)programming means propagation of the FC transcriptional identity to all syncytial nuclei via auto-activation of *col* transcription [15, 16]. In both *col* ^*ΔLCRM1.3*^ and *col* ^*ΔLCRM0.5*^ embryos, Col is present in PCs at stage 11 and undetected in syncytial DA3 nuclei at stage 15 (Figure 3A), indicating that complete *versus* incomplete DA3 transformation results from different Col dynamics during the fusion process. At stage 14, Col is indeed detected in muscle precursors in *col* ^*ΔLCRM0.5*^, but not *col* ^*ΔLCRM1.3*^ embryos (Figure 3B). To trace the origin of this difference, we analysed in more detail *col* transcription in the DA3 PC, FC and stage 14 syncytium by FISH/immunostaining using hemizygous embryos which display one hybridisation dot per active nucleus (Figure 3C). In wt embryos, a dot is systematically detected in the DA3/DO5 PC (20/20 segments; 5 embryos analysed), the DA3 FC (20/20) and 80% of DA3 nuclei at stage 14 (6 of 7-8 nuclei per fibre on average; 27 segments). A *col* dot is always detected as well in the DA3/DO5 PC in *col* ^*ΔLCRM1.3*^ (21/21) and *col* ^*ΔLCRM0.5*^ (18/18) embryos, reflecting *col*-ECRM activity. In *col* ^*ΔLCRM1.3*^embryos, it is only detected in a minor fraction of FCs (4/15), while completely lost at stage 14 (0/6-7 nuclei per fibre on average; 24 segments). In *col* ^*ΔLCRM0.5*^ embryos, however, it is maintained in the DA3 FC (19/19) and still detected at stage 14, in one nucleus, likely the FC nucleus (21 segments), sometimes two nuclei. The duration of *col* transcription correlates with the detection of Col protein. Taken together, the col ^ΔLCRM1.3^ and col ^ΔLCRM^0.5 expression data and muscle identity phenotypes show that iTF transcription in the FC is strictly required for muscle identity. However, acquisition of a specific morphology also requires identity reprogramming of “naïve” syncytial nuclei (Figure 3D).

**Figure 3.**
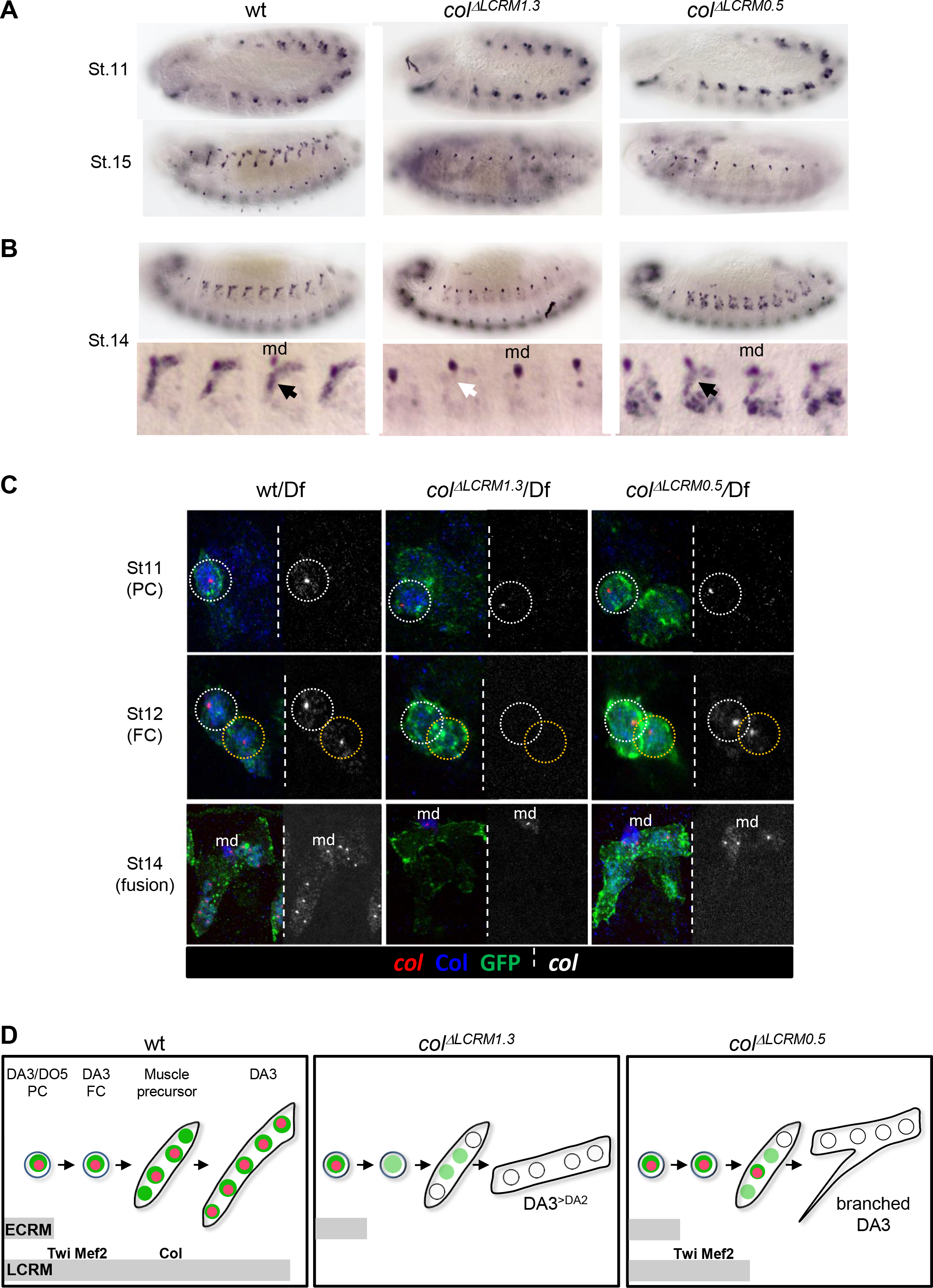
Identity reprogramming of syncytial nuclei controls the final muscle morphology. **(A)** Col immunostaining of wt, *col* ^*ΔLCRM1.3*^and *col* ^*ΔECRM0.5*^ embryos, showing a normal pattern at stage 11 and complete absence of Col protein at stage 15 in *col* ^*ΔLCRM*^ embryos. **(B)** Immunostaining stage14 embryos with a close up view of 4 segments where position of the md neuron is indicated. It shows the presence of low amounts of Col protein (black arrow) in the growing DA3 muscle in *col* ^*ΔECRM0.5*^, and absence in *col* ^*ΔLCRM1.3*^embryos (white arrow). **(C)** *col* transcription (red dots), Col protein (blue) and *col*-LCRM-moeGFP expression (green) in the DA3/DO5 PC, stage 11, DA3 FC, stage 12, and developing DA3 muscle, stage 14, in *wt*, *col* ^*ΔLCRM1.3*^ and *col* _*ΔECRM0.5*_/*Df(2L)^BSC429^*; *col*-LCRM-moeGFP embryos. In each panel, Col protein (right) is shown separately in black and white. *col* transcription ceases at the PC stage in *col* ^*ΔLCRM1.3*^ embryos and does not propagate to other syncytial nuclei in *col* ^*ΔLCM0.5*^ embryos. **(D)** Schematic representation of the dynamics of *col* transcription (red dots) and nuclear Col (green) in the DA3/DO5 PC, the DA3 FC, muscle precursor, and DA3, DA3^>DA2^ and branched DA3^>DA3/DA2^ muscles in wt, *col* ^*ΔLCRM1.3*^ and *col* ^*ΔECRM0.5*^ embryos, respectively. Temporal activity of *col*-ECRM and *col*-LCRMs is represented by horizontal grey bars.

### Muscle transformation and branched muscles; attraction and repulsion

Co-staining of stage 16 embryos for *col*-LCRM-moeGFP and integrin confirms that the DA3 posterior edge anchors to dorsal tendon cells and anterior edge to lateral intersegmental tendon cells, giving it its final acute shape [1] (Figure 4A). The formation of DA3^>DA2^ and branched DA3^>DA3/DA2^ muscles in *col* ^*ΔLCRM*^ embryos recalls transient attachment of the wt DA3 muscle to anterior dorsal tendon cells along the anterior segmental border, giving it a characteristic angled-shape at stage 14 [24]. Filopodial contacts with these cells are still observed at stage 17 but they exclusively reach either edge of the DA3 muscle attachment site (MAS) from the adjacent posterior segment (Figure 4A). This suggests that they are attracted by tendon cells and repelled from the DA3 MAS. In contrast, in *col* ^*ΔLCRM1.3*^embryos, the MAS surfaces of adjacent DA3^>DA2^ muscles partly or completely align with each other over the intersegmental border. In some segments, the anterior DA3 edge maintains a MAS with lateral tendon cells, resulting in DA3^>DA3/DA2^ branched muscles. Phalloidin staining of ready-to-hatch wt embryos confirms that the posterior DA3 and DA2 anterior MASs precisely align with each other, forming a heterotypic DA3/DA2 MAS. On the contrary, in *col* ^*ΔLCRM1.3*^embryos, homotypic MASs between adjacent DA3^>DA2^ muscles are privileged. As a consequence, the DA2 anterior MAS is shifted dorsally and the staggered rows architecture of DA muscles is lost (Figure 4B). Of note, only the internal muscles show precise MAS matching, while intermediate muscles such as DO1, DO2 and DO4 muscles show shifted MAS matching (Figure 4B).

**Figure 4.**
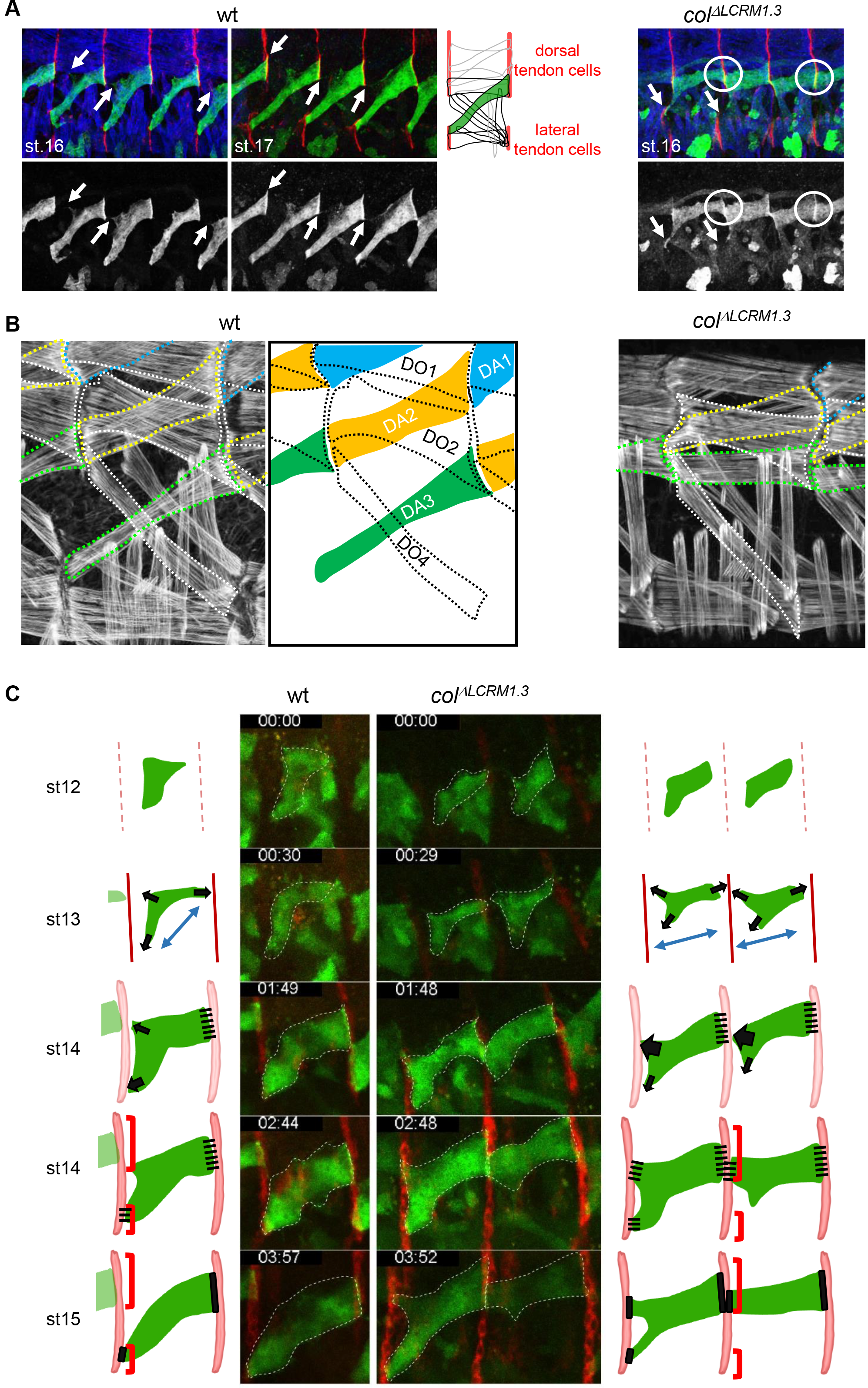
Imaging of aberrant DA3 muscle attachment in *col*-LCRM mutants. **(A)** Immunostaining of stage 16 and 17 wt and stage 16 *col* ^*ΔLCRM1.3*^embryos for integrin (red), *col*-LCRM-moeGFP (green) and Myosin Heavy chain (blue, stage 16 only). moeGFP alone is shown below in black and white; white arrows point to filopodial projections reaching the edges of the posterior DA3 MAS in wt embryos and lateral tendon cells in *col* ^*ΔLCRM1.3*^embryos, with homotypic DA3^>DA2^ MASs being circled. A schematic diagram of the dorsal and dorso-lateral muscles, and the dorsal and lateral tendon cells positions (red) in stage 17 wt embryos is shown. **(B)** Phalloidin staining of stage 17 wt and 16 *col* ^*ΔLCRM1.3*^embryos. Close up views of the DA3 and DA3^>DA2^ MASs are shown, with DA1 circled in blue, DA2 circled in yellow and DA3 or DA3^>DA2^ circled in green. A drawing illustrates wt muscle matching, with DA1, DA2 and DA3 muscles coloured in blue, yellow and green respectively, and DO muscles circled by dotted lines. **(C)** Live imaging of moeGFP expressed in the DA3 lineage under control of and tendon cell precursors labelled by Stripe-Gal4 × UAS-redFP. Live embryos were filmed during 4 hours (Sup. Movies 1 and 2) and Z sections collected every ~2-2,5 minutes. The outlines of the developing muscles are depicted schematically for each stage. In both wild-type (left, one segment shown), and *col* ^*ΔLCRM1.3*^embryos (right, two segments shown), the posterior muscle end reaches the intersegmental border first. In stage 14 wt embryos, DA3 contacts both dorsal and lateral tendon cells (red brackets), along the anterior border. The dorsal projections then retract and MAS maturation gives the DA3 muscle its final acute orientation, stage 15. In *col* ^*ΔLCRM1.3*^embryos, the anterior dorsal projections fail to retract, leading to homotypic DA3^>DA2^ attachment.

Live imaging of lateral-oblique and ventral transverse muscles development distinguished 3 phases of muscle elongation [25]: 1) FC migration, stage 12, ending with the first FC/FCM fusion event and stretching of the muscle precursor along an axis; 2) bipolar myotube elongation, characterised by the presence of extensive filopodia at both axis ends, in search for attachment sites, stages 13 to 15; 3) maturation of myotendinous attachment, stage 16. To explore further how heterotypic *versus* homotypic MASs form, we live-imaged wt and *col* ^*ΔLCRM1.3*^ embryos, starting at stage 12 (defined as t_0_ in **Movies 1 and 2**). The DA3 muscle was labelled by *col*-LCRM-moeGFP and tendon cell precursors along the entire intersegmental border by *stripe* Gal4xUASmCD8RFP [26]. Live-imaging of wt embryos (**Movie 1;** Figure 4C) shows that the DA3 muscle precursor extends protrusions directed towards dorsal tendon cells, posteriorly and lateral tendon cells, anteriorly, conform to a bipolar extension scheme [25]. However, numerous filopodia emanating from its dorsal surface are also observed, which extend toward anterior dorsal tendon cells, giving the DA3 muscle precursor a peculiar triangular shape (stage 12). At stage 13, the posterior DA3 edge contacts dorsal tendon cells, while the anterior edge(s) reach the intersegmental border slightly later, a gap also observed during live-imaging of a ventro-lateral muscle [27]. This transiently results in three contact sites, early stage 14. At late stage 14, the surface contact between the DA3 posterior edge and dorsal tendon cells spreads while a stable MAS form. The anterior ventral connection in turn becomes stable, while dorsal anterior filipodia now retract, as they are repelled from the DA3 posterior MAS. In support of this mechanism of homotypic repulsion, remaining filopodia only contact the rims of DA3 anchoring (Figure 4A). In *col* ^*ΔLCRM1.3*^ embryos, the main axis of the DA3^>DA2^ muscle precursor elongates more longitudinally than wt, already at stage 12 (**Movie 2;** Figure 4C). At stage 13, filopodia emanating from the dorsal surface of the DA3 precursor are more active than ventral filopodia, suggesting that repulsion from dorsal tendon cells is lower than wt. Moreover, there is no repulsion/retraction of filopodia contacting dorsal tendon cells, along the anterior border where the DA3^>DA2^ muscle of the adjacent segment anchors. Rather, stabilization of these projections leads to formation of the abnormal DA3^>DA2^ homotypic MASs seen by integrin staining of fixed embryos (Figure 4A). Projections to lateral tendon cells are sometimes also stabilized, giving rise to DA3^>DA3/DA2^ branched muscles. In conclusion, live imaging of wt embryos shows that the final acute orientation of the DA3 muscle results from first attachment of the posterior edge to dorsal tendon cells, repulsion of the anterior edge from this MAS and its stable attachment to lateral segmental tendon cells. Homotypic repulsion is alleviated upon DA3^>DA2^ identity shift, leading to formation of abnormal, homotypic MASs.

### Muscle identity shifts provoke mismatching of muscle attachments

The viability and DA3>DA2 phenotype of *col* ^ΔLCRM^ larvae provided the opportunity to assess the impact of embryonic muscle patterning defects on *Drosophila* larval crawling. We first recorded the fraction of DA3, DA3>DA2 and branched muscles in 3^rd^ instar wt and *col* ^*ΔLCRM*^ larvae, by live imaging individual larvae expressing the muscle-specific isoform of Mhc fused to GFP, MHC-GFP (Figure 5A). It conforms to the statistics made in late embryos, except for an increased proportion of branched muscles, which could reflect under-evaluation of their proportion in embryos, due to threshold detection limit of *col*-LCRM-moeGFP expression in thin fibers (Figure 5B). Since DA muscles are internal and form indirect MASs, we then examined the pattern of muscle/muscle attachments, using scanning electron microscopy (SEM) of dissected larval filets (Figure 5C; Figure 5 Sup1). In wt larvae, and as seen in embryos (Figure 4), the anterior edge of the DA3 muscle anchors to a nodal lateral attachment site shared with the LL1 muscle while its posterior edge, spreads more and matches the DA2 muscle in the next posterior segment. A heterotypic alignment is also seen for the DA2/DA1 MASs. Thus, staggered ends of DA1, DA2, DA3 muscles both draw strength lines spanning 3 adjacent segments, and create regular tension surfaces between segments (Figure 5C; Figure 5 Sup1–Sup2). In *col*^*ΔLCRM*^ larvae, a DA3^>DA2^ /DA3^>DA2^ homotypic alignment substitutes for the DA3-DA2 heterotypic alignment and the DA3 muscle is no more connected to the nodal lateral attachment site (Figure 5C). As a result, the DA1-DA2-DA3 strength line is broken. An additional consequence of DA3^>DA2^ homotypic attachment is loss of the regular staggered ends architecture of the DA3-DA2-DA1 alignment (Figure 5 Sup1). In case of DA3^>DA3/DA2^ branched muscles, a DA3-LL1 MAS is maintained, although of reduced surface, and two strength lines co-exist (Figure 5C and Figure 5 Sup1). In conclusion, SEM analyses of larval muscles show that loss of DA3 identity leads to abnormal, homotypic MASs involving several dorsal and dorso-lateral internal muscles and distortion of DA muscle strength lines and staggered ends architecture.

**Figure 5.**
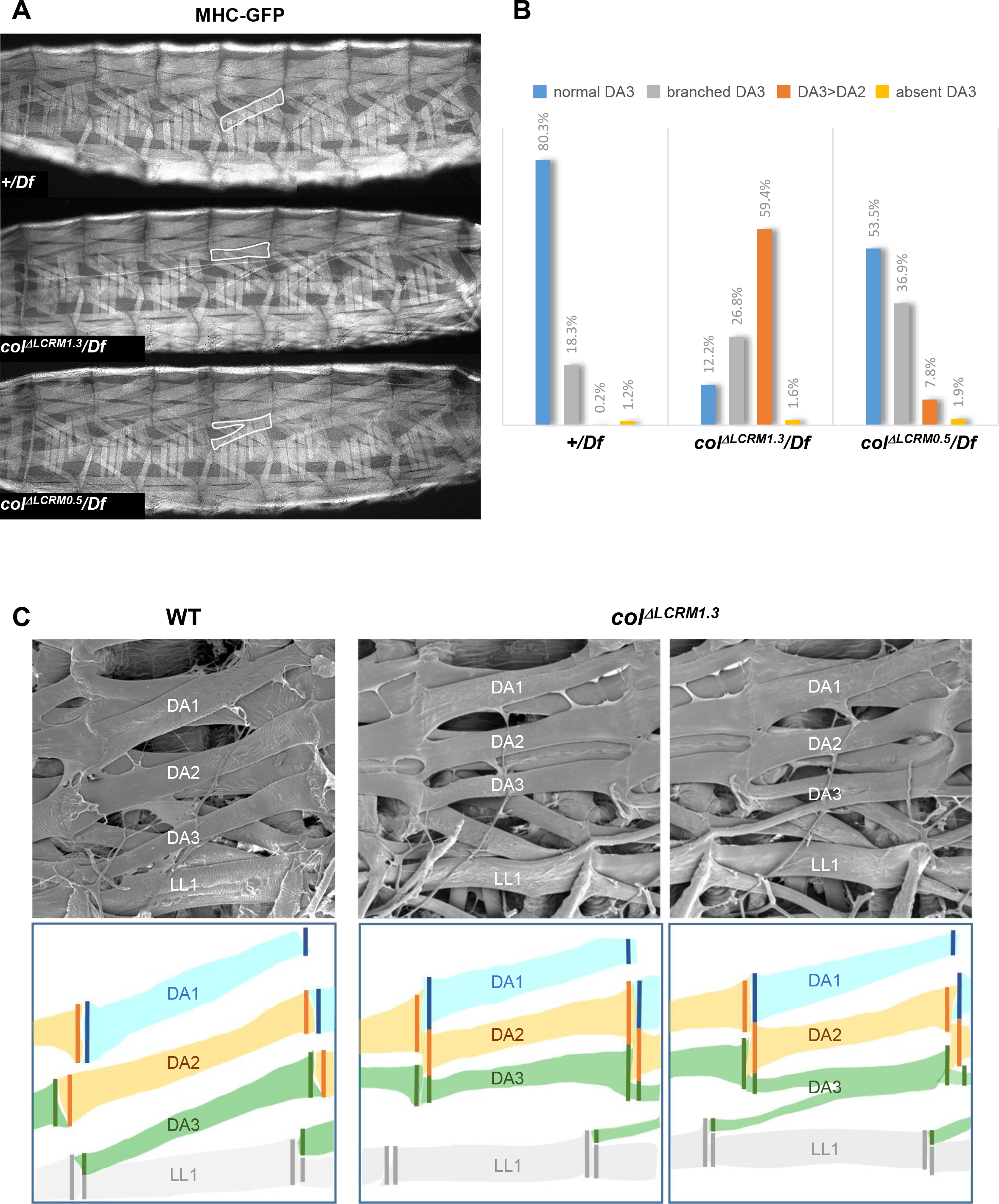
Disrupted muscles matching in *col*-LCRM mutant larvae. Live-imaging of the somatic musculature of wt, *col* ^*ΔLCRM1.3*^ and *col* _*ΔECRM0.5*_/*Df(2L)^BSC429^;* MHC-GFP 3^rd^ instar larvae; lateral views, anterior to the left. Examples of DA3, DA3^>DA2^ and DA3^>DA3/DA2^ muscles are circled in dotted white. **(B)** Quantification of the relative proportions of DA3, branched DA3^>DA3/DA2^, DA3^>DA2^ and absence of DA3 muscles in *wt*, *col* ^*ΔLCRM1.3*^ and *col* ^*ΔECRM0.5*^/*Df(2L)^BSC429^;* MHC-GFP larvae. Segments A1 to A7 have been considered. *(+/Df*, n=375 segments/27 larvae; *col*^*ΔLCRM1.3*^/*Df* n=320/23 larvae; *col*^*ΔLCRM0.5*^/*Df*, n=264/19). **(C)** Scanning electron microscopy of filleted larvae showing the dorso and dorso-lateral internal muscles. Below is a schematic color-coded drawing. In wt larvae, the DA1, DA2 and DA3 muscles are parallel to each other in each segment with precise matching of the DA3/D2 and DA2/DA1 MASs at each posterior segmental border. In *col*^*ΔLCRM1.3*^larvae, DA3^>DA2^ muscles show homotypic MASs over dorsal tendon cells and the DA3/LL1 connection is lost. The posterior MASs of branched DA3^>DA3/DA2^ is composite. Formation of homotypic DA3^>DA3/DA2^ MASs.

### Branched muscles result in specific locomotion defects

Drosophila larval crawling relies upon ordered abdominal body wall muscle contractions [28]. The ectopic muscle tendon and muscle-muscle attachments observed in *col*^*ΔLCRM*^ larvae raised the question of its possible impact on locomotion. To address this question, we compared the locomotion of *+/Df(2L)^BSC429^, col^Δ^LCRM1.3^^/Df(2L)^BSC429^* and *col ^Δ^LCRM0.5^^/Df(2L)^BSC429^* larvae using FIM (FTIR-based Imaging Method) and the FIMTrack software, which allows tracking many larvae simultaneously and quantitatively describing a variety of stereotypic movements [29–31]. We focused on selected parameters: crawling speed, stride length, stride duration. Other parameters such as head cast duration, angular speed and amplitude, were recorded but not further considered here. We first recorded the “walking rate”, also called crawling speed, during the first 20 seconds (5 frames/second), after larvae have been dropped on the agarose gel. At that time, larvae engage an “escape response” corresponding to an active crawling phase (Figure 6A). Box-plot graphs (left) show a large intra-variability for each of the 3 genotypes. Beyond this variability, we observe, however, that *col ^Δ^LCRM0.5^^/Df^BSC429^* larvae display a significantly reduced crawling speed on average 1.05±0.034 mm/sec (n=108), compared to 1.15±0.031 mm/sec for *+/Df(2L)^BSC429^* controls (n=118), (p=0.03) (Figure 6A). To further investigate the origin of this speed reduction, we measured two crawling speed parameters: stride length and stride duration (Figure 6B-C). A significantly shorter stride was measured for *col* ^*Δ^LCRM0.5^*^/*Df(2L)*^*BSC429*^ larvae (1.08±0.025 mm) compared to control *+/Df(2L)^BSC429^* larvae (1.17±0.024 mm), (p=0.008). Furthermore, stride duration was extended, from (1.09±0.014 sec) for *+/Df(2L)^BSC429^* larvae to 1.15±0.017 sec for *col ^Δ^LCRM0.5^^/Df(2L)^BSC429^* larvae. For both crawling speed, stride length and stride duration, *col*^*ΔLCRM1.3*^/*Df(2L)*^*BSC429*^ larvae (n=112) display intermediate values. Yet, differences with either control or *col ^Δ^LCRM0.5^^/Df(2L)^BSC429^* larvae fall below the significance threshold level. All together, the data show a significantly decreased crawling speed of *col ^Δ^LCRM0.5^^/Df(2L)^BSC429^* larvae, corresponding to a shorter stride length coupled with increased stride duration.

**Figure 6.**
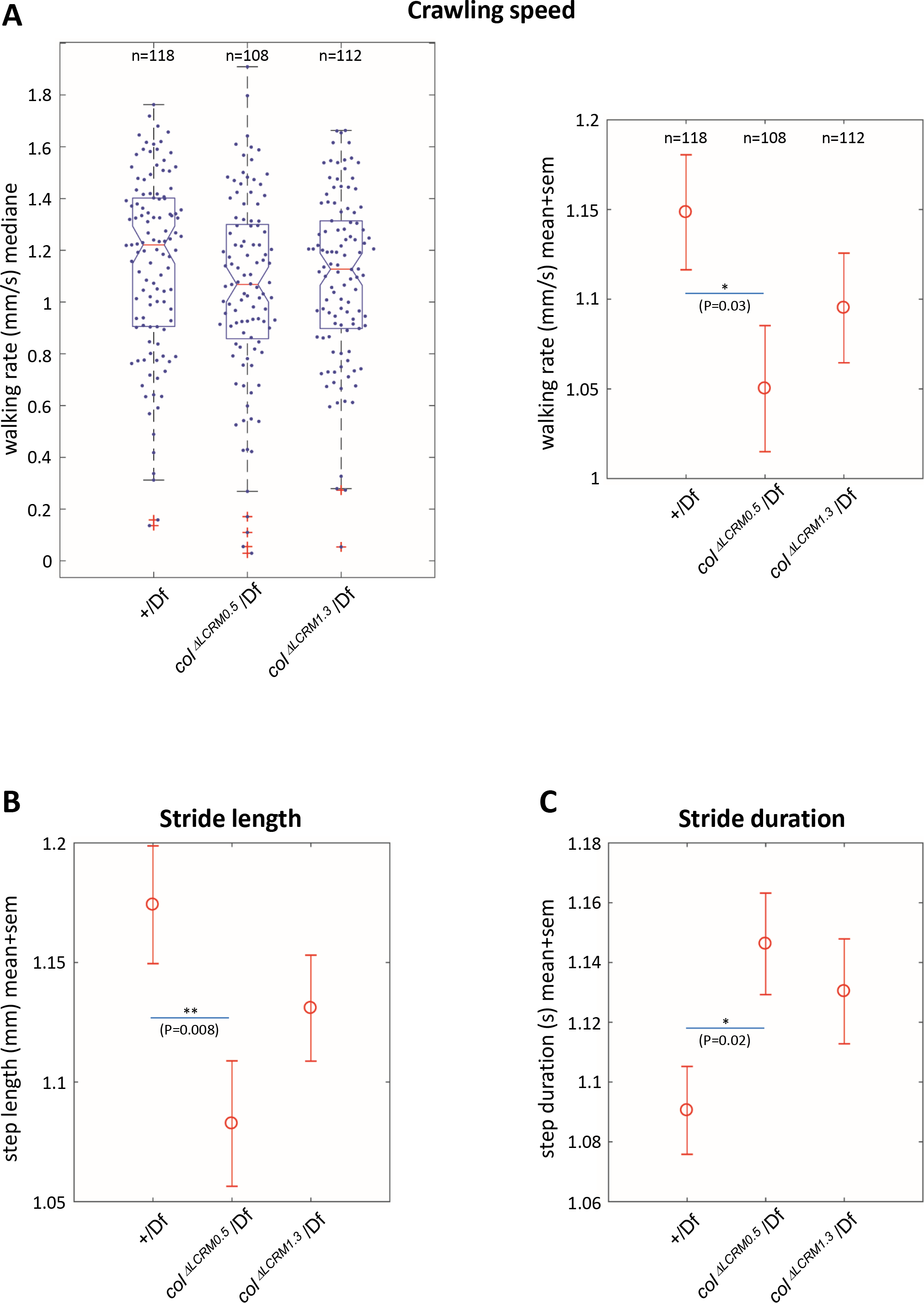
Branched muscles result in specific locomotion defects. **(A)Left**, Tukey’s diagram (box-plot graph) showing the walking rate in mm/s (crawling speed) of +/*Df*, *col* ^*ΔECRM0.5*^/*Df* and *col*^*ΔLCRM1.3*^/*Df* larvae. For each genotype, one point represents the average measurements for one larva, recorded during 20 seconds. 50% of the points are located within the Tukey’s diagram. The red line represents the median and the narrowed area represents the confidence interval of the median (95%). **Right**, same data as left, showing the mean speed ± standard error of mean (SEM) for each genotypes. **(B)** Stride length in mm. **(C)** stride duration in s. The number of larvae (n) tested for each genotype is indicated in **(A)**. Only the significant differences are indicated (* p <0.05 and ** p <0.01).

## Discussion

The stereotyped set of 30 somatic muscles in each abdominal segment which underlies larval crawling has been thoroughly described many years ago [1, 32]. One essential aspect laid out at that time was the concept of founder cell (FC), *i.e.*, the assignment to form a distinctive muscle to a single founder myoblast able to recruit other myoblasts by fusion. The morphological identity of each muscle is foreseen by a specific pattern of iTF expression in its FCs [33, 34]. Here, we report the setting of a CRM deletion strategy connecting iTF expression to the musculature architecture and larval locomotion.

### iTF transcription in muscle PCs; redundant CRMs

Some iTFs are only transiently transcribed during *Drosophila* muscle development, *e.g*., *Kr*, and *nau*, others such as *col*, *ladybird* and *S59*, at every step of the process [10, 13, 15, 16, 35, 36]. Reporter analyses indicated that *col* transcription was controlled by two separate, sequentially acting CRMs, of overlapping activity at the PC step, suggesting a handover mechanism between positional and maintenance CRMs at this stage [24]. However, CRM deletion analyses show that *col*-LCRM activity in specific muscle PCs is not dependent, but redundant with *col*-ECRM activity, and separate from autoregulatory *col*-LCRM activity that is specific to the DA3 FC and syncytial nuclei. This leads to a model where iTF code refinement at each step of muscle identity specification, PMC>PC, PC>FC and FC>syncytial nuclei, is driven by a separate CRM. Distribution of the PC-specific information into two CRMs further supports the idea that iTF regulation at the PC stage is nodal to muscle identity specification [7, 10, 24, 36–38]. Of note, the present analysis concentrated on the DA3 lineage where *col* expression is maintained, whereas *col* is expressed in several PCs (Figure 1 Sup1). Morphological transformations of other dorso-lateral muscles are observed at variable frequency in *col* protein null mutants [9], suggesting that *col*-ECRM activity provides robustness to the combinatorial control of identity of these muscles. PC/muscle specific CRMs have only been functionally identified for a handful of muscle iTFs [39]. Computational predictions identified, however, several thousand putative muscle enhancers and uncovered extensive heterogeneity among the combinations of transcription factor binding sites in validated enhancers, beside the core intrinsic muscle regulators Tin, Mef2 and Twi [17, 40, 41]. Dissecting whether step-specific and redundant/distributed CRM configurations [19, 42, 43] apply to many muscle iTFs and underlie the progressively refined control of the final muscle pattern, is a future challenge.

### Distinctive muscle morphology requires identity reprogramming of fused myoblasts

The process by which iTFs determine the final morphological features of each muscle fiber, is not fully understood. Fusing FCM nuclei generally adopt the iTF protein code of the FC nucleus, while propagation of iTF transcription and activation of realisation genes is lineage-specific [11–15, 44]. We have previously shown that Col protein import precedes activation of *col* transcription in fused FCM nuclei, and correlates with activation of realisation genes, a sequence of events termed syncytial identity reprogramming [15, 16]. Upon loss of *col* transcription in the DA3 FC, there is a complete DA3^>DA2^ transformation. When *col* transcription is not propagated from the FC to syncytial nuclei there is an incomplete DA3^>DA3/DA2^ transformation. From this, two conclusions can be drawn: 1) Final selection of myotendinous connection sites is intrinsic to FC identity. 2) Identity reprogramming of syncytial nuclei is required for robustness of this selection.

### Identity shifts and branched muscles

The formation of DA3^>DA3/DA2^ branched muscles recalls the transient attachment of the wt DA3 muscle to three groups of tendon cells, a process deviating from the bipolar migration/attachment scheme and little documented, so far [3, 24, 25, 45]. It could be specific of muscles which, like DA3 and unlike DA2, bind different groups of tendon cells, dorsal, lateral and ventral, at their anterior and posterior ends (Figure 4). Interestingly, “asymmetrical” attachment is conferred to the DA2 muscle by Col ectopic expression, as observed in *tup* mutant embryos [11]. This suggests that asymmetric selection is an active, highly regulated process. A few molecules involved in targeted attachment of subsets of muscles to specific tendon cells have been characterized: Kon tiki/Perdido, a single pass transmembrane protein and the PDZ protein DGrip which are required for proper elongation of ventral longitudinal muscles [46]; the ArfGAp protein Git is required for sensing integrin signaling and halting elongation of Lateral Transverse (LT) muscles once their attachment site has been reached [45, 47]; Robo/Slit signaling attracts muscles at segmental borders, the Slit ligand being expressed by tendons, and Robo and Robo2 receptors by elongating muscles. *slit* also acts as a short-range repellent contributing to the collapse of leading-edge filopodia when a muscle reaches the tendon extracellular matrix [48, 49]. *In vivo* imaging showed that the DA3 muscle fails to stop at the segment border in *slit* mutants and sometimes branches [49]. However, the DA3 *slit* branching pattern is different from that observed in *col* ^*ΔLCRM*^ mutants, and rather similar to that frequently observed in *nau* mutants, with two posterior attachment sites in place of one [10, 11]. Of note, some *Drosophila* larval muscles are normally branched, such as DA1, which dorsally connects two different sets of tendon cells and the DO1 dorsal-oblique muscle (Figure 5 Sup2) [50]. Which genes downstream of iTFs are responsible for asymmetric attachment site, *i.e*., proper balancing attraction/repulsion cues, remains unknown and should be the focus of future studies.

The stabilisation of branched muscles in *Drosophila* identity mutants is one important finding as branched muscle fibers are observed in humans, following muscle regeneration after damage or in muscular dystrophy pathologies [51]. It should allow for the first time to investigate, *in vivo*, the functional properties of branched muscles and associated mechanical instability in an otherwise normal muscle pattern.

### Muscle staggered ends; heterophilic *versus* homophilic interactions

In addition to tendon cells, internal lateral muscles adhere to other muscles [4, 32] resulting in a very precise configuration of contraction force lines. The DA3/DA2 and DA2/DA1 MASs precisely match over each segmental border, such that the DA1, DA2, and DA3 perfectly align over 3 consecutive segments Figure 4B). Live imaging DA3 and DA3^>DA2^ development shows that the DA3/DA2 matching in wt embryos results from homotypic repulsion: first connection of the posterior tip of DA3 to dorsal tendon cells leads to repulsion of filopodia issued from the same muscle in the next segment and heterotypic DA3/DA2 MASs. In parallel, the DA3 anterior edge is attracted to lateral tendon cells. Upon DA3^>DA2^ identity shift, homotypic repulsion does not occur and homotypic DA3^>DA2^ MAS are stabilized. Interestingly, this does not exclude DA3^>DA2^/DA2 matching (Figures 4 and 5). Cell matching is a widely used process during embryogenesis to construct complex tissue architecture. Selective filopodia adhesion has recently been shown to ensure precise matching between identical cell types and boundaries between different cell types in the *Drosophila* heart. In this case, homotypic matching is linked to differential expression by each cell type of the adhesion molecules, Fasciclin III and Ten-m [52]. The generation of different muscle-specific identity mutants should, in the next future, allow us to determine whether and how heterochrony between posterior and anterior adhesion, differential expression of attractive or repulsive molecules, and/or competition between muscles for the same tendon cells contributes to precise staggered ends of internal muscles.

### Branched muscles impact on crawling speed

*Drosophila* larval crawling is a well-suited paradigm to link muscle contraction patterns and locomotor behaviour. Longitudinal, acute and oblique muscles within a larval segment contract together and, as they begin to relax, the contraction is propagated to the next segment, creating a peristaltic wave from tail to head (forward locomotion), or from head to tail (backward locomotion) [28]. In many organisms, the rhythmic movements of locomotion are part of behavorial routines that facilitate the exploration of an environment. Exploratory routines mostly alternate straight line movement also called “active crawling phase”, with change of direction and acquisition of a new trajectory, the “reorientation phase” [53–55]. The active larval crawling phase requires an intense, prolonged muscular effort. In contrast, during the reorientation phase, larvae remain at the same spot while bending and moving their head, often followed by a turning event. In this study, we focused on crawling parameters during the active phase. The crawling speed during the escape response was significantly reduced in *col ^Δ^LCRM0.5^^/Df^BSC429^*, compared to control *+/Df(2L)^BSC429^* larvae, whereas *col^Δ^LCRM1.3^^/Df(2L)^BSC429^* larvae showed an intermediary, not significant phenotype. This could seem paradoxical because the number of DA3^>DA2^ transformed muscles is higher in *col^Δ^LCRM1.3^^/Df(2L)^BSC429^* than *col ^Δ^LCRM0.5^^/Df(2L)^BSC429^* larvae while *col ^Δ^LCRM0.5^^/Df(2L)^BSC429^* larvae present more branched muscles. These observations indicate that i) single muscle transformations only moderately impact crawling speed, raising the possibility of biomechanical compensation by other muscles; ii) branched muscles could be less efficient than fully transformed muscles. At this point, the reason why, – either mechanic weakness, improper innervation or Ca^2+^ wave propagation, antagonistic force lines upon muscle contraction, and/or possible induced bilateral asymmetry - may only be object of speculation. The present Drosophila data provide a novel entry point to studying the pregnant question of how the physiological properties of branched fibres which progressively accumulate in Duchenne muscular dystrophy patients differ from morphologically normal fibres.

## Material and Methods

### Fly strains

All *Drosophila melanogaster* stocks and genetic crosses were grown using standard medium at 25°C. The strains used were *white^[1118]^, col*^Late^CRM 4-0.9 [9], *col^1^* [12], *sr-Gal4*/TM6 (obtained from G. Morata, Madrid, Spain). The 12 *kn* Janelia-Gal4 lines (GMR) [56], *UAS-mcd8RFP, Mhc-GFP*, *Df(2L)^BSC429^*, *kn*^*MI15480*^ and *vasa-cas9*^*VK00027*^ lines were provided by Bloomington Drosophila Stock Center. The *col^1^* and Df(2L)BSC429 strains were balanced using CyO,{*wg*^*en11*^-lacZ} or CyO,{*dfd-YFP*} and homozygous mutant embryos or larvae identified by absence of lacZ or YFP expression, respectively.

### CRM deletions generated by Crispr/Cas9

Genomic col target sites were identified using http://tools.flycrispr.molbio.wisc.edu/targetFinder/ [57]. Prior to final selection of RNA guides (gRNA) for deletions of *col* CRMs, genomic PCR and sequencing of DNA from *kn*^*MI15480*^ and *vasa-cas9^VK00027^* flies was performed to check for polymorphisms in the targeted regions. Guides targeting *col*-ECRM and *col*-LCRM were inserted in the pCFD4: U6:3-gRNA vector (Addgene n^o^: 49411) as described [58]; (see http://www.crisprflydesign.org/wp-content/uploads/2014/06/Cloning-with-pCFD4.pdf). All guides were verified by sequencing. The sequences of the oligonucleotides used to construct each gRNA expression plasmid are given in Supplemental Material, Figure ure 1 Sup2 and 3. To delete the core region of *col*-LCRM*, vasa-cas9* embryos were microinjected with gRNAs in pCFD4 (200 ng/μl). To delete the *col*-ECRM, *kn*^*MI15480*^ embryos were injected with gRNA in pCFD4 (150 ng/μl) and pAct-Cas9U6 (400 ng/μl). Each adult hatched from an injected embryo was crossed to the balancer stock *sna*^*Sco*^/CyO, {*wg*^*en11*^-*LacZ*} and 100-200 F1 fly were individually tested for either *col* CRM deletion by PCR on genomic DNA.

### Reporter constructs, immunohistochemistry, in situ hybridization

The *yellow* intron (*yi*), FlyBase ID #FBgn0004034 (position: 356918-359616) was inserted in the *lacZ* coding region between aa (Tyr 952) and aa (Ser 953) by standard PCR-based cloning position. Antibody staining and *in situ* hybridization with intronic probes were as described previously [16]. Primary antibodies were: mouse anti-Col (1/50; [16]), mouse anti-LacZ (1/1000; Promega), mouse anti-βPS (1/10; Developmental Studies Hybridoma Bank), rabbit anti-GFP (1/1000; Torrey Pines Biolabs), rabbit anti-β3-tubulin (1/5000; from R. Renkawitz-Pohl, Marburg, Germany); chicken anti-GFP (1/500; Abcam), Phalloidine-Texas RedX (1/500; Thermofisher Scientific). Secondary antibodies were: Alexa Fluor 488-, 555- and 647- conjugated antibodies (1/300; Molecular Probes) and biotinylated goat anti-mouse (1/2000; Vector Laboratories). Digoxygenin-labelled antisense RNA probes were transcribed *in vitro* from PCR-amplified DNA sequences, using T7 polymerase (Roche Digoxigenin labelling Kit).

*In situ* Hybridization with Stellaris RNA FISH probes were done as described by the manufacturer for *Drosophila* embryos (https://www.biosearchtech.com). The FISH probe sets for *col* were designed using the Stellaris probe designer (https://www.biosearchtech.com/stellarisdesigner) and labeled with a Quasar 670 Dye (Stellaris Biosearch Technologies). One set of 48 oligonucleotides was designed against the first *col* intron to detect primary nuclear transcripts. Another was designed against exons to detect cytoplamic mature transcripts. Both sets recognize all *col* RNA isoforms and were used together. When antibody staining and FISH were combined, the standard immuno-histochemistry protocol was performed first, with 1U/l of RNase inhibitor from Promega included in all solutions, followed by the FISH protocol. Confocal sections were acquired on Leica SP8 or SPE microscopes at 40x or 63x magnifications, 1024/1024 pixel resolution. Images were assembled using ImageJ and Photoshop softwares.

### Phenotype quantification at embryonic and larval stages

To quantify embryonic phenotypes, *col*-LCRM-moeGFP embryos were immunostained with a primary mouse anti-GFP (1/500) (Roche) and secondary biotinylated goat anti-mouse (1/2000) (VECTASTAIN^®^ ABC Kit). Stained embryos were imaged using a Nikon eclipse 80i microscope and a Nikon digital camera DXM 1200C. A minimum 100 A1-A7 abdominal segments of stage 15-16 embryos were analyzed for each genotype. (+/Df: n=127 segments - 16 embryos; *col*^*ΔECRM*^/Df: n=170 segments - 23 embryos; *col*^*ΔLCRM1.3*^/Df: n=190 segments - 27 embryos; *col*^*ΔLCRM0.5*^/Df: n=103 segments - 13 embryos). To quantify larval phenotypes, wandering L3 larva displaying *MHC-GFP* reporter line were immobilized between slide and coverslip, and left and right larval sides imaged using Nikon AZ100 Macroscope at 5x magnification. Minimum 260 abdominal segments were analyzed for each genotype. (+/Df: n=375 segments - 27 larvae; *col*^*ΔLCRM1.3*^/Df: n=320 segments - 23 larvae; *col*^*ΔLCRM0.5*^/Df: n=264 segments - 19 larvae).

### Live imaging embryonic muscle development

Embryos were bleach dechorionated and stage 12 embryos manually picked, laterally orientated and mounted on a coverslip coated with heptane glue to prevent drift during imaging. A drop of water was placed on the embryos to maintain their survival. Images were collected on a Leica TCS-SP8 confocal using a 25X water immersion lens. Sections were recorded every 130 to 150 seconds for the wt embryos and every 120 to 160 seconds for the *col*^*ΔLCRM1.3*^ embryos, and z-stacks collected with optical sections at maximum 1 µm interval. Image processing was performed with Fiji (http://fiji.sc/wiki/index.php/Fiji) and custom programming scripts in Fiji. The z-stacks projections were corrected in x and y dimensions by manual registration using a reference point tracking.

### Scanning Electron Microscopy (SEM)

To prepare fillets, third instar wild type and homozygous *col*^*ΔLCRM1.3*^ larvae raised at 25°C were dissected in myorelaxant buffer, according to [59]. Larvae were cut longitudinally on the ventral side to preserve and expose the dorsal and dorso-lateral musculature. Fillets were then fixed 1 hour in a 4% formaldehyde/ 2.5% glutaraldehyde mixture in 1X PBS, washed in water and dehydrated gradually in ethanol. Fillets were dried at the critical point (Leica EM CPD 300 critical point apparatus), covered with a platinum layer (Leica EM MED 020 metalliser) and imaged with a Quanta 250 FEG FEI scanning microscope.

### Behavioral analysis

We conducted locomotion assays by tracking the trajectory of larvae using the FIM method [29]. Wandering third instar larvae were gently picked up with a paintbrush and transferred to an agar plate. The larvae were then videotaped using a digital camera (Baumer VCXG53M); lentille (Kowa LM16HC); infrared filter (IF093SH35.5) and tracked using the FIMTrack software [30]. Each video containing 5 to 10 larvae per run, on a 1% agarose gel, was recorded at 5 frames/sec for 20 seconds. Analysis was done by using MATLAB software.

## Acknowledgements

We thank the Bloomington Stock Center for *Drosophila* strains. We gracefully thank Cristian Pasquaretta (CRCA) for his help in statistical analyses of locomotion behaviour, Patrick Arrufat (CRCA), Philippe Firmin and Claude Nexon (CBD) for their assistance in building the locomotion tracking table, Brice Ronsin, Toulouse RIO Imaging platform and Julien Favier, *Drosophila* embryos microinjection platform.

---

**Figure 1 Sup1:**
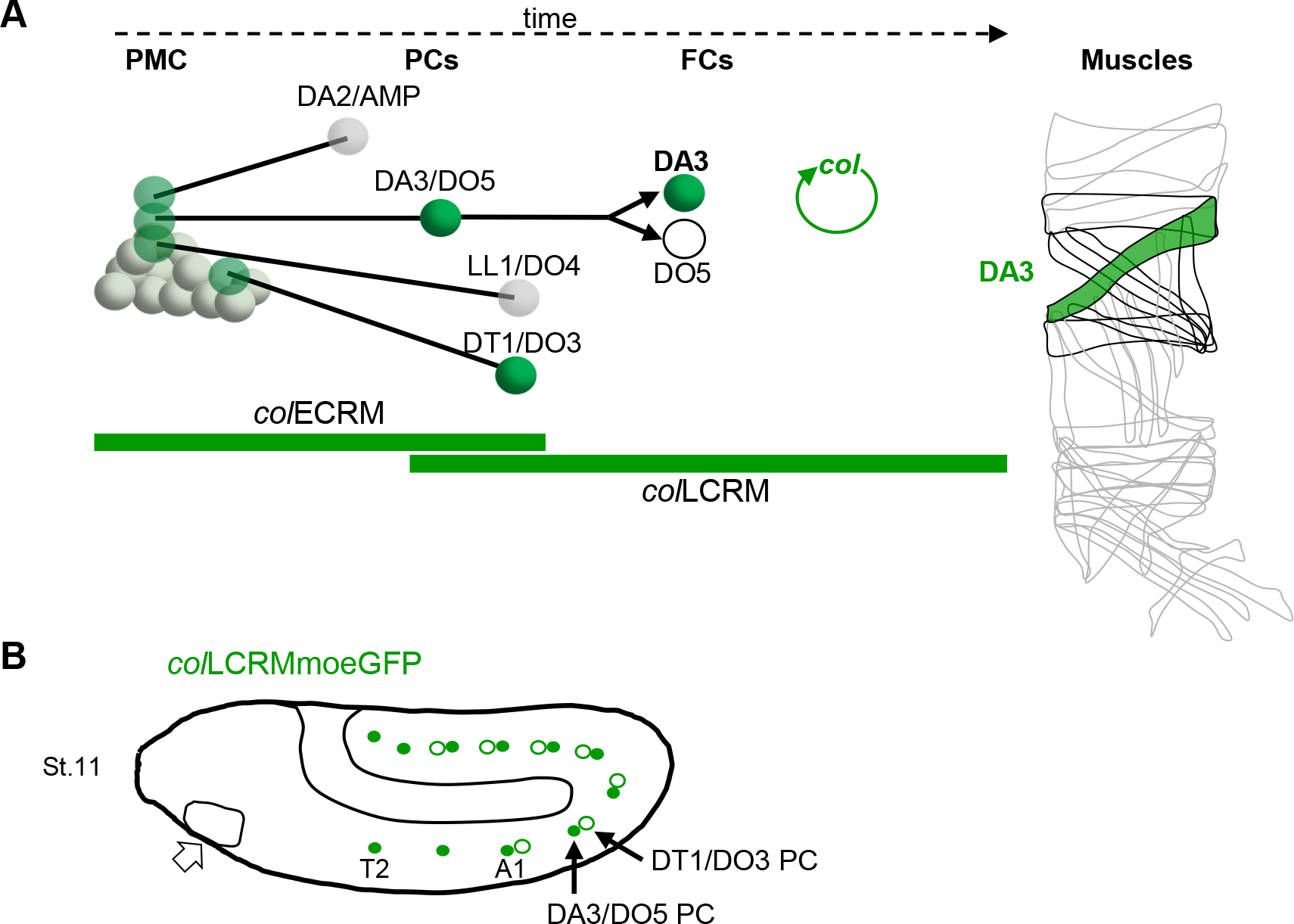
Summary of Col expression in muscle development. **(A)** Diagrammatic representation of the sequential emergence of 4 different PCs (green) from the Col-expressing PMC (grey) and division of the DA3/DO5 PC into 2 FCs and the *col* auto-regulatory loop operating in the DA3 lineage; the names of each PC and FC are indicated. Accumulation of Col protein is in green; the time windows of mesodermal *col*-ECRM and *col*-LCRM activity are indicated by green bars. Right, muscle pattern of an abdominal segment highlighting the DA3 muscle (green). The other muscles derived from PCs transiently expressing Col are indicated by thicker contours. **(B)** Schematic representation of *col*-LCRMmoeGFP expression in a stage 11 wt embryo. The head expression domain is indicated by an open arrow, the DA3/DO5 PC (green, T2 to A9 segments) and DT1/DO4 PCs (circled green, A1-A7 segments) by black arrows in A2.

**Figure 1 Sup2.**
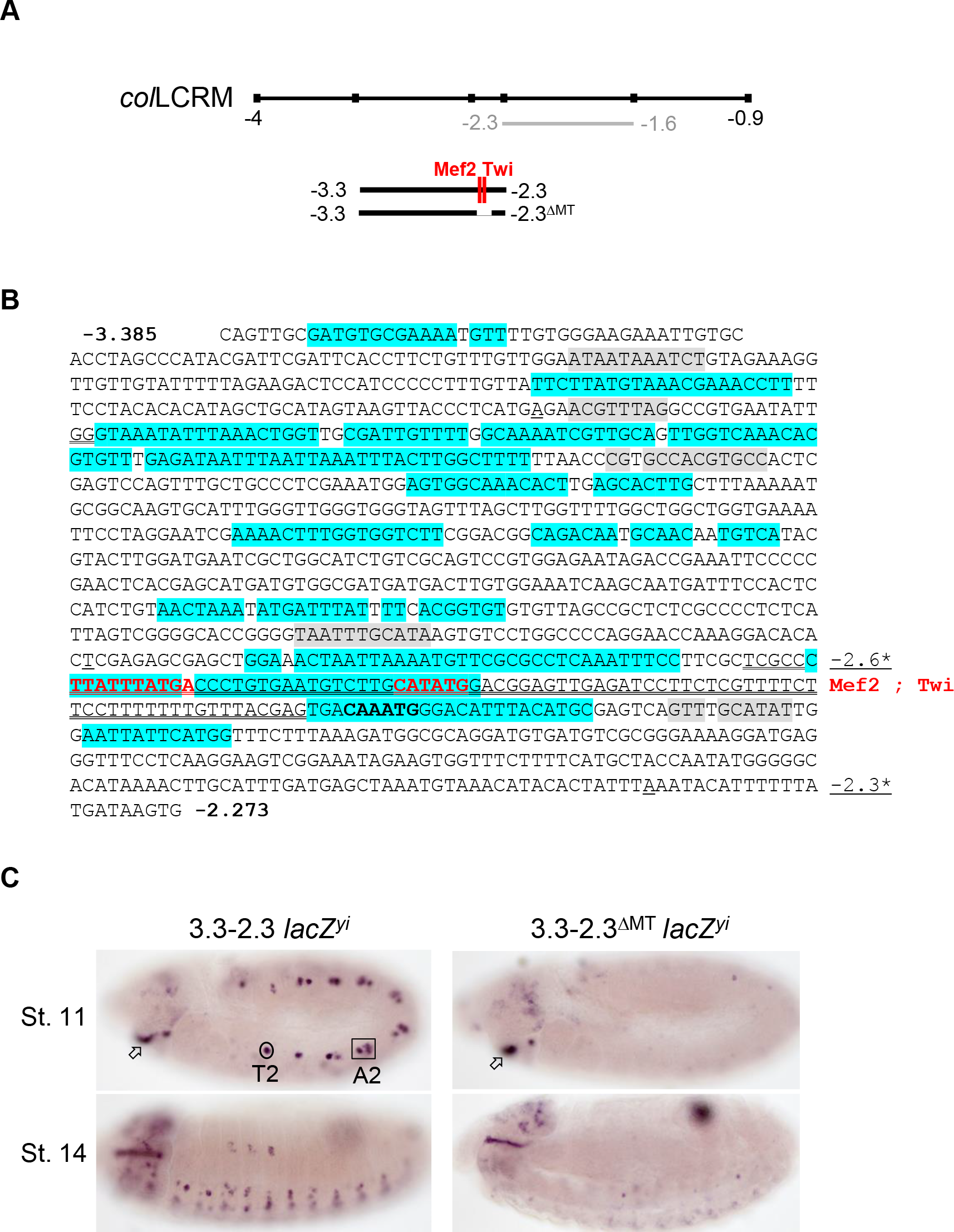
Mapping of a *col* PC-specific CRM. **(A)** Schematic representation of *col*-LCRM (see Figure 1) indicating the positions of the −2.3 to −1.6 fragment (grey) previously shown to drive expression in the DA3 syncytium [16] and new reporter constructs, −3.3-2.3 and −3.3-2.3 ^ΔMT^ with the position of conserved Mef2 and Twi binding sites. **(B)** Nucleotide sequence of the −3.3-2.3 fragment, positions −3385 to −2273. The endpoints of previously characterised reporter fragments indicated **\***in the margin [9, 16] are underlined. The −20-bp sequence motifs that are more than 90% conserved at the same relative positions in *D. melanogaster*, *D. pseudobscura* and *D. virilis* are shaded in blue, additional shorter conserved motifs, shaded in grey. Binding sites for Mef2 and Twi (red) are in bold. The 97bp segment containing both Mef2 and Twi binding and deleted in 3.3-2.3^ΔMT^ is underlined. **(C)** *In situ* hybridisation to primary reporter transcripts (*yellow* intron (*yi*) inserted in the *lacZ* coding sequence). The 3.3-2.3 fragment drives specific expression in the DA3/DO5 and DT1/DO4 PCs (circled in segment T2 and framed in A2) and the intercalary segment (arrow). Its activity in PCs is lost upon deletion of the region encompassing Mef2 and Twi binding sites (3.3-2.3^ΔMT^ reporter).

**Figure 2 Sup1:**
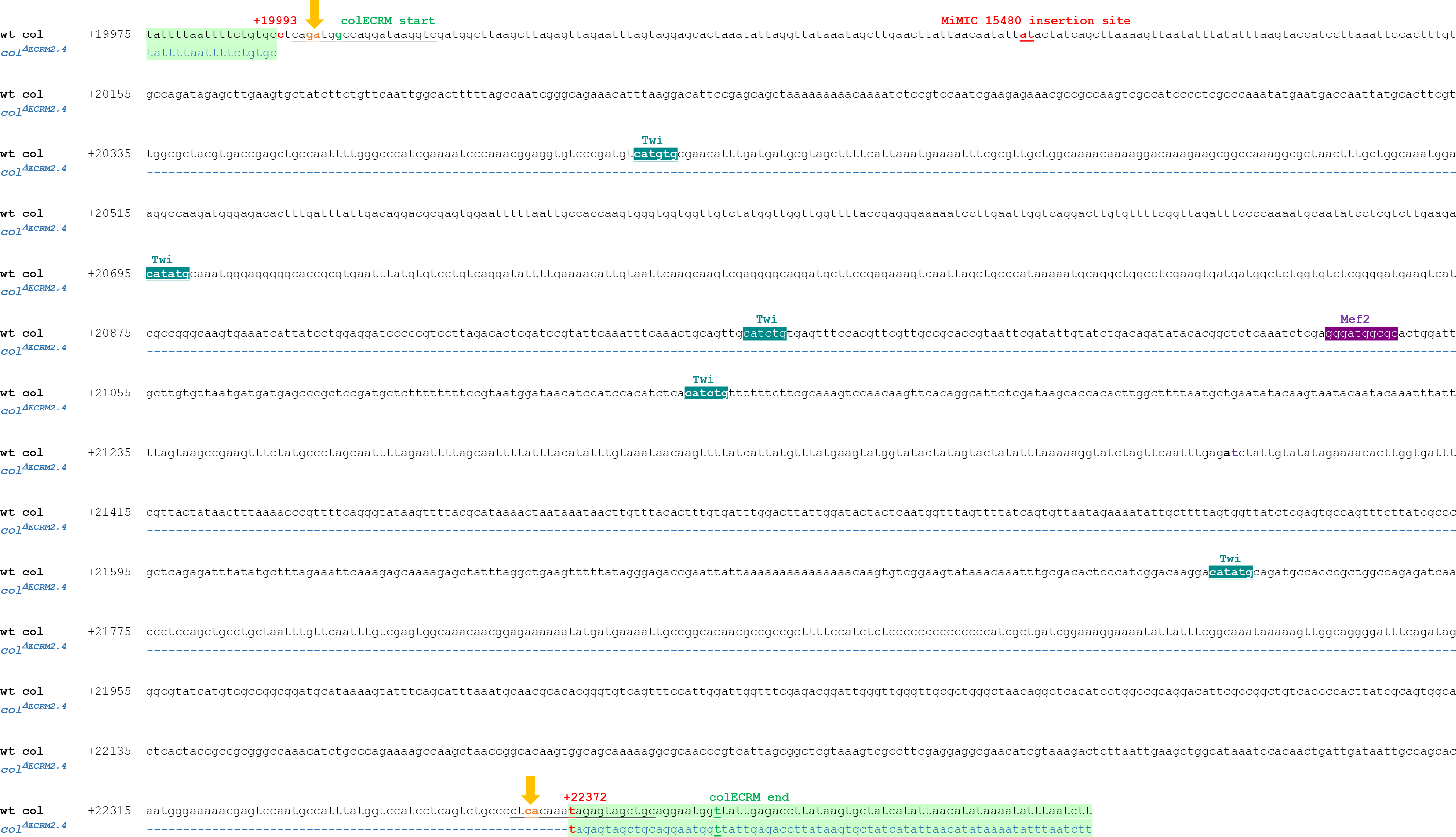
*col*-ECRM deletion. Genomic sequence spanning the previously mapped *col*-ECRM [9] (start and end indicated in bold, green) and the *col* ^*ΔECRM2.4*^ deleted fragment (dashed blue line). The nucleotide positions of the *col* ^*ΔECRM2.4*^ deletion endpoints (bold red) are given relative to the *col* transcription start site. The predicted cleavage sites of the used sgRNAs (sequences underlined) are indicated by yellow arrows. The MiMIC 15480 is inserted between the red underlined nucleotides. Predicted binding sites for Twist (blue) and Mef2 (purple), conserved between *Drosophila* species.

**Figure 2 Sup2:**
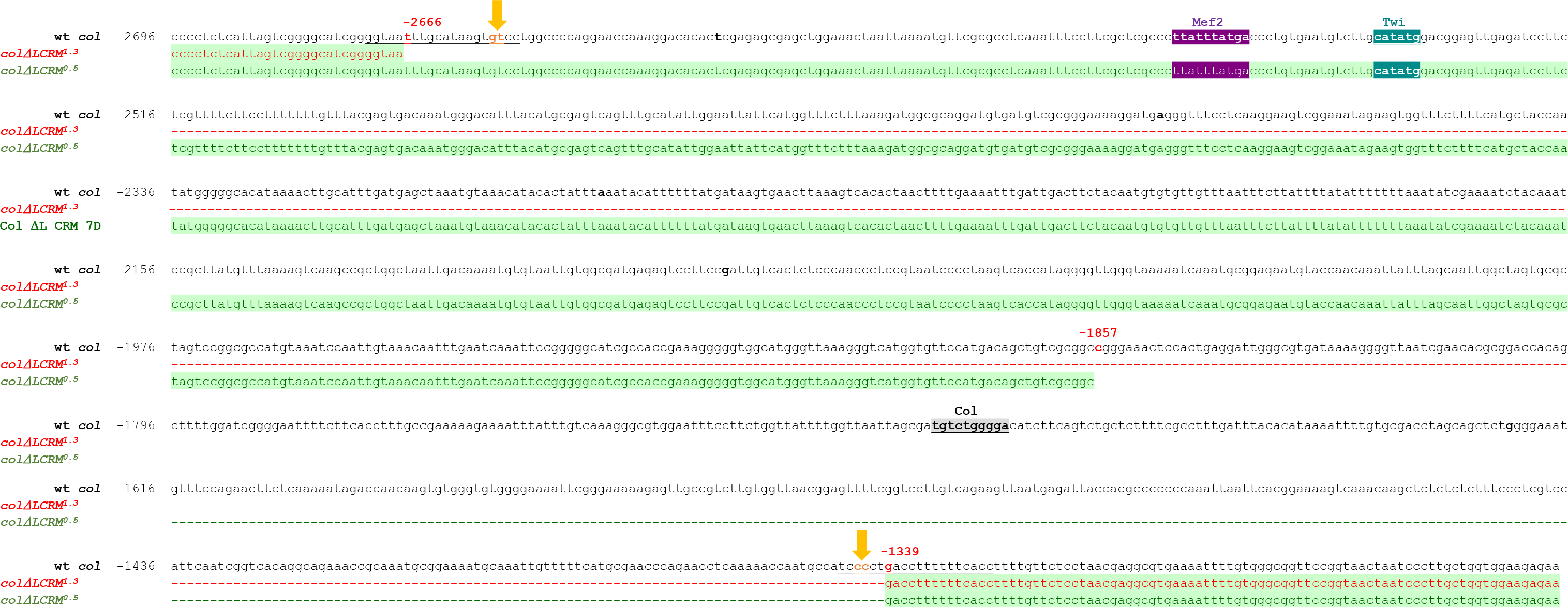
*col*-LCRM deletions. Genomic sequence spanning the *col* ^*ΔLCRM1.3*^(from −2666 to −1339) and *col* ^*ΔLCRM0.5*^ (from −1857 to −1339) deletions. Deleted fragments are represented by dashed lines and their coordinates given relative to *col* transcription start site. The predicted cleavage sites of the used sgRNAs (sequences underlined) are indicated by yellow arrows. Predicted binding sites for Twist (blue), Mef2 (purple), and Col (grey), conserved between *Drosophila* species.

**Figure 5 Sup1.**
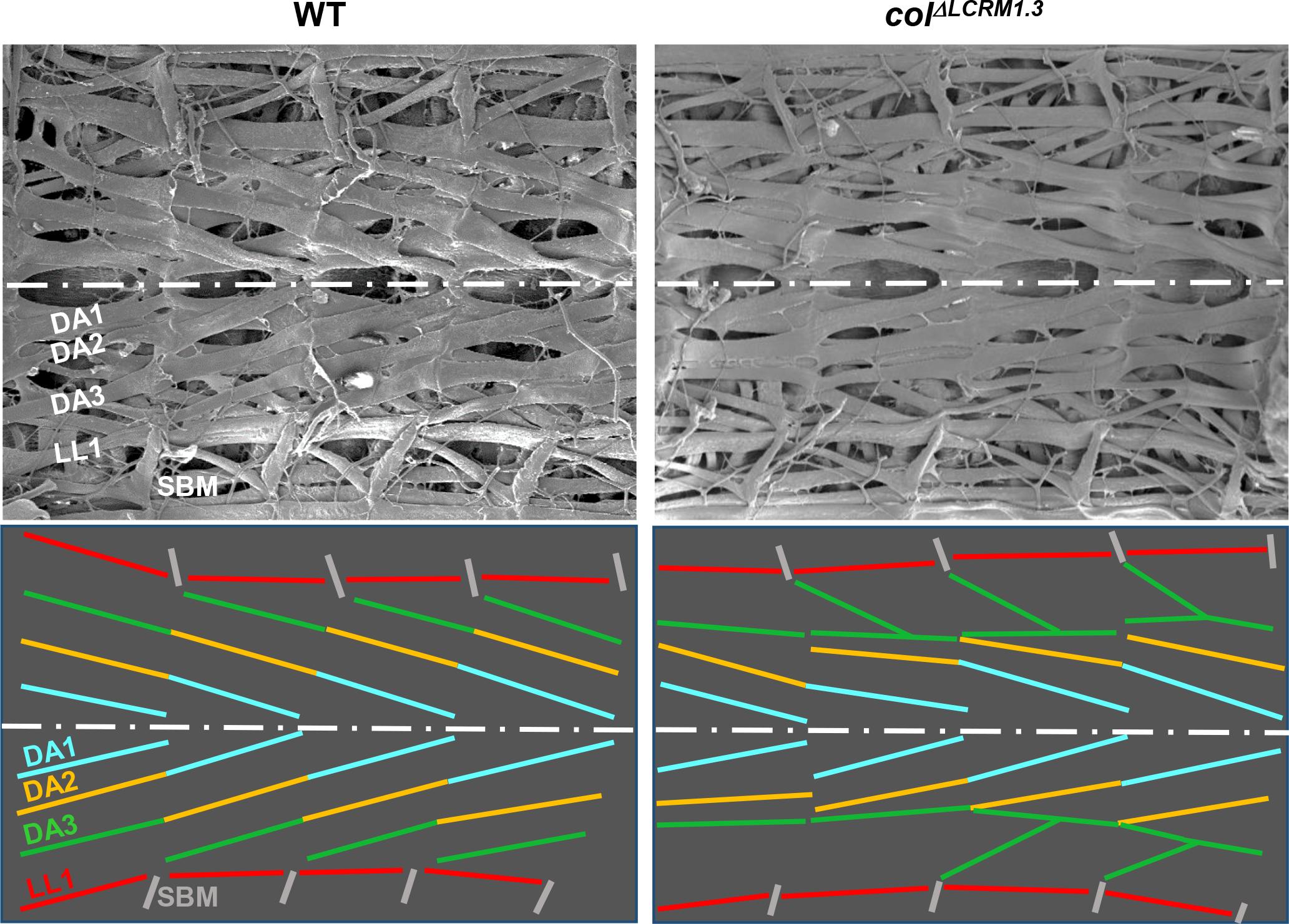
Scanning electron microscopy of wt and *col* ^*ΔLCRM1.3*^ internal muscles. Larval fillets cut longitudinally along the ventral midline to expose the dorsal and dorso-lateral internal muscles. The dorsal midline is represented by a white dashed line. Upper panels: 4 abdominal segments of wt and *col* ^*ΔLCRM1.3*^L3 larvae. Lower panels: muscle orientations are schematized by colored lines, DA1 yellow, DA2 blue, DA3 green, LL1 red and SBM grey. In a wild type larva, the DA1, DA2 and DA3 muscles are oriented parallel and draw a force line over 3 consecutive segments and straight tension lines between two adjacent segments. In *col*^*ΔLCRM1.3*^larvae, DA3^>DA2^ and DA3^>DA3/DA2^ transformations these lines are broken due to the formation of homotypic DA3^>DA2^ MAS and either loss or reduction of the DA3/LL1 connection.

**Figure 5 Sup2.**
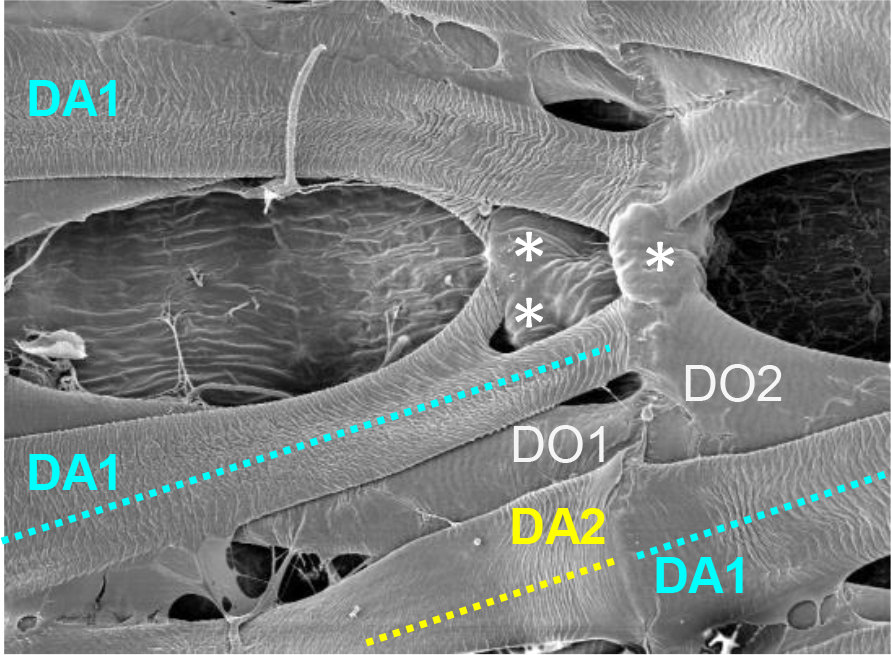
Scanning electron microscopy of dorsal muscle attachment. Larval fillets were prepared as in Figure ure 5 Sup1. The dorsal DA1 attachment sites are shown. On its dorsal posterior side, one DA2 branch connects an internal tendon cell (blue asterisk) and the DO2 muscle. The second, dorsal-most branch connects to a separate, superficial tendon cell (white asterisk). On its anterior ventral side, DA1 matches DA2 from the adjacent anterior segment.

